# Unveiling the dominant role of soil pH in shaping nitrogen cycling microbial gene abundances: Insights from 65-years of chemical fertilizer selection in acidic grassland meadow

**DOI:** 10.1101/2024.04.26.591135

**Authors:** Akari Mitsuta, Kesia Silva Lourenco, Mart Ros, Yoshitaka Uchida, Eiko Eurya Kuramae

## Abstract

Understanding the microbial processes driving the nitrogen (N) cycle is crucial for enhancing plant productivity and mitigating environmental pollution. The long-term application of synthetic fertilizers induces significant alterations in the microbial community functions. Microbes inhabiting acidic soils may exhibit distinct responses to chemical fertilizer application compared to soils with neutral pH. However, the chronic or occasional changes resulting from repeated nutrient flushes due to fertilizer application remain insufficiently elucidated, especially in acidic grasslands. Therefore, our study was on an acidic semi-natural grassland, where the soil was subjected to chemical fertilizer P (superphosphate), K (potassium sulfate), PK, N (ammonium nitrate), NPK, PK+N (PK applied in spring and N applied once in summer) over a span of 65 years. Gene abundances associated with the N-cycle (*amoA*, *nifH*, *nirK*, *nirS*, and *nosZ*) were quantified at seven different time points throughout the year considering the temporal effect caused by fertilizer application. Our findings reveal that in the long term, soil pH emerged as the predominant factor influencing the gene abundance related to N-fixation and denitrification outweighing the impact of nutrient availability. Notably, the application of N fertilizer had a positive effect on the abundance of nitrifiers, while the abundance of denitrifiers decreased due to soil acidification induced by fertilizer application. In summary, our study highlights that the microbial community involved in N cycling is more sensitive to the difference in soil pH shaped by long-term fertilizer application rather than to the direct impact of fertilizer application.

**Highlight:**

- Seasonal sampling confirmed the long-term effect of fertilizer on microbial N-cycling genes
- Fertilizer types shaped soil pH, consequently impacted on diazotrophs and denitrifies abundances
- Abundance of denitrifying microbe did not explain the increase of N_2_O emission by N fertilizer
- N_2_O emission was positively correlated with AOA abundance

## Introduction

To meet the increasing demand for food, there has been a significant intensification of agricultural systems and a continuous rise in the consumption of chemical fertilizers, including nitrogen (N), phosphorus (P), and potassium (K) fertilizers over the last few decades (FAO, 2019). While intensive fertilization has led to a substantial increase in food production, the sustainability of this practice is being questioned due to potential negative environmental and ecological consequences associated with the long-term and repeated use of synthetic fertilizers. One major concern is the shift of soil microbial function related to the N cycle, as soil microorganisms play a crucial role in the transformation of N, involving vital processes such as N-fixation, nitrification, and denitrification (Kuypers et al., 2018). This microbial activity constitutes a pivotal aspect for sustaining the agricultural system because microbial N transformation is a key process in increasing plant productivity and reducing environmental pollution such as the emission of N_2_O, a gas with a global warming potential 273 times greater than that of carbon dioxide (CO_2_) (Forster et al., 2021).

The application of chemical fertilizer, including N, P, and K, is a strong driver of microbial diversity and activity related to N-cycling, however, the effect of fertilizer treatment could be dependent on the combination of the element and duration of the application. For instance, long-term N fertilizer application enhances the abundance of nitrifiers and denitrifiers (Chu et al., 2007; Zhou et al., 2015), whereas a negative impact on diazotrophs abundance by N fertilizer application is reported (Wang et al., 2017, 2022). However, when N fertilizer is accompanied by P and K, the impact of fertilizer on diazotrophs abundance can result in less negative than a single application of N (Hu et al., 2019; Wang et al., 2017). Furthermore, the duration of the fertilizer application was indicated as an important factor influencing the abundance of microbial N-cycling genes. A meta-analysis indicates that the effect size of N fertilizer on the abundance of *amoA*-AOB, *nirS*, *nirK*, and *nosZ* genes differs between soils receiving mineral fertilizer for more than 20 years and those receiving it for less than 20 years (Ouyang et al., 2018). However, since fertilizer application causes a temporal nutrient flush and a shift in the microbial community, seasonal sampling is essential to evaluate the long-term effects of fertilizer application. Thus, systematic sampling along with fertilizer application in different seasons would help in achieving a comprehensive understanding of how each element of fertilizer (P, K, and N) results in the dynamics of the functional microbial community associated with N cycling.

In addition to nutrient availability, soil pH is a ubiquitous factor that strongly shapes the microbial community. Microorganisms inhabiting acidic soil may respond differently to the application of chemical fertilizers than microbes living in soils with neutral pH. Considering the effect of soil pH on the microbial N cycle is therefore crucial, as pH is one of the most important factors determining the growth of microbes (Fernández-Calviño and Bååth, 2010). For instance, the application of N into acidic soil (pH < 5) resulted in a negative influence on microbial biomass but showed no significant effect on the abundance of denitrifying and nitrifying microbial communities (Geisseler and Scow, 2014; Ouyang et al., 2018). On the other hand, at higher pH, N application has a positive influence on microbial biomass, denitrifiers, and nitrifiers abundance. Notably, the application of N fertilizer is one of the common reasons for soil acidification in agricultural lands (Tian and Niu, 2015), and the shift in soil pH by N fertilizer often determine as the most explanatory factor for the abundance and community of N cycling microbes (Chen et al., 2015; He et al., 2007; Wang et al., 2017). Given the considerable potential for expanding the world’s agricultural land, acidic soil, which encompasses approximately 40% of the Earth’s surface, emerges as a more critical area for crop production (Bian et al., 2013). However, information is still lacking, on the interplay between nutrient inputs and soil pH in acid soils that have been conditioned by repeated fertilizer applications and how it shapes the microbial N-cycling community.

The Ossekampen long-term grassland experiment was established in 1958 in Wageningen, the Netherlands, and represents an acidic semi-natural grassland. Over 65 years, the grassland has consistently received various chemical fertilizers, including P, K, PK, N, NPK, and PK+N (with PK applied in spring and N in summer). Due to the acidity of the soil in this grassland (pH around 4.5 in the plot without fertilizer), the outcomes were expected to be different from similar experiments conducted in grassland with a neutral pH. Previous research has been conducted to explore the long-term impact of fertilizer application on plant and microbial abundance, community, and diversity (Cassman et al., 2016). However, the long-term effect on the abundance of N-cycling microbes has not been thoroughly quantified. Therefore, this study aimed to investigate shifts in the microbial N cycling community, focusing on ammonia oxidation (*amoA*), denitrification (*nirS*, *nirK*, *nosZ*), and N-fixation (*nifH*), with quantitative real-time PCR (qPCR). The primary objective was to comprehend the impact of long-term synthetic fertilizer application on the abundance of microbial N-cycling genes in the acidic temperate grassland. We hypothesize that 1) the chronic effects of long-term synthetic fertilizer application significantly shape the N-cycle microbial community, and 2) the observed shift in gene abundance is unique to acidic soil compared to soil with neutral pH found in the literature.

## Materials and Methods

### Site description

The Ossekampen long-term grassland experiment (51 degrees 58′15″N; 5 degrees 38′18″E, Wageningen, The Netherlands) was started in 1958 in an extensively grazed species-rich grassland with soil characterized as heavy river clay. Prior to the experiment, the land was grazed and had been used in alternate years for haymaking. The treatments consist of control (no fertilizer application), P (superphosphate), K (potassium sulfate), PK (combination application of P and K), N (ammonium nitrate), NPK (combination application of N, P, and K), PK+N (application of PK in spring and single application of N in summer). Fertilizer was applied April 11 and July 12 and the amount of applied fertilizer was described in Table 1. The grass is mown twice yearly, in July and October before fertilization, and the biomass is removed from a 2.5-m-wide perimeter to prevent the dispersal of seeds into other plots. The plots are duplicate 40-m^2^ (16 m× 2.5 m) plots for each fertilizer treatment and treatments are separated by unfertilized 2.5-m-wide buffer strips to avoid contamination (dispersal of seeds) between treatments, and these strips are similarly maintained by mowing.

**Table 1.** Amount of applied fertilizer.

| Treatment | Fertilizer application (kg/ha) |  |  |  |  |  |
| --- | --- | --- | --- | --- | --- | --- |
|  | First application |  |  | Second application |  |  |
|  | N | P | K | N | P | K |
| Control | 0 | 0 | 0 | 0 | 0 | 0 |
| P | 0 | 22 | 0 | 0 | 0 | 0 |
| K | 0 | 0 | 108 | 0 | 0 | 0 |
| PK | 0 | 22 | 165 | 0 | 0 | 0 |
| N | 100 | 0 | 0 | 60 | 0 | 0 |
| NPK | 100 | 33 | 311 | 60 | 0 | 0 |
| PK+N | 0 | 33 | 311 | 160 | 0 | 0 |

### Soil sampling

Soil samples (150 g) were collected from the 0–10 cm top layer in 2022. For both soil and gas sampling, each plot was split into two sub-plots and duplicated. Sampling was conducted three times in spring (April 6, 13, and 19) and four times in summer (July 7, 14, 18, and August 2). Thus, a total of 168 samples (6 treatments × 4 replications × 7 time points) were collected. Immediately after sampling, soil subsamples (30 g) were stored at −20 °C until the molecular and chemical analyses.

### DNA extraction and quantification of N-cycle gene abundance

Total soil DNA was extracted from 0.40 g of soil using the MoBio PowerSoil DNA Isolation Kit (MoBio, Solana Beach, CA, USA) according to the manufacturer’s instructions. DNA quantity and quality were determined using a NanoDrop ND-1000 spectrophotometer (NanoDrop Technologies, DE, USA). The abundances of the functional genes *amoA*-AOB, *amoA*-AOA, *amoA*-comammox, *nirS* clusters I-IV, *nirK* clusters I-IV, *nosZ* clades I and II, and *nifH* and ribosomal RNA genes for total bacteria, archaea, and fungi were quantified by qPCR using the CFX96 Touch™ Real-Time PCR Detection System (Bio-Rad). The qPCR standard curve of each functional gene was constructed as described in the supplemental material. For each primer set, the cycling conditions, annealing temperature, cycle number, and primer amount were optimized, and the amplicon of the target gene was checked with electrophoresis (Table S1). The genes *nirS* clusters II-IV and *nirK* clusters III and IV were not detected in our samples.

### Soil chemical property analysis

Soil mineral N (NH_4_⁺₋N, NO_2_ ^−^₋N, and NO_3_ ^−^₋N) was measured with a continuous flow analytical system (Skalar San^++^ continuous flow meter) after extraction with 2 M KCl in 1:5 (soil: solution). Soil pH was measured in 1:2.5 in 0.01 M CaCl_2_. Available P-phosphate was determined by calorimetry, and exchangeable cations (K^+^, Ca^2+^, and Mg^+^) were determined by atomic absorption spectrometry (Shimadzu AA-7000) analysis of the soil extract obtained using ion-exchange resin (van Raij et al., 2001). Cationic micronutrients [iron (Fe), manganese (Mn), copper (Cu), and zinc (Zn)] were extracted in a solution (pH 7.3) containing 0.005 M diethylenetriaminepentaacetic acid (DTPA), 0.1 M triethanolamine (TEA), and 0.01 M CaCl_2_ and determined by atomic absorption spectrometry. Total C and N from soils were analyzed using a CN elemental analyzer (Perkin Elmer 2400).

### Greenhouse gas measurement

In parallel with soil sampling, CO_2_ and N_2_O fluxes were measured using a PICARRO G2508 ring-down spectroscopy gas analyzer (Picarro Inc., Santa Clara, CA, USA). These fluxes were measured with a closed chamber technique (Clough et al., 2020). One week before the measurement, polyvinylchloride (PVC) rings with an inner diameter of 19 cm and a height of 10 cm were inserted 5 cm into the soil, at two locations per plot, at 1.5 m from both edges of each plot. For the flux measurements, a white opaque polypropylene flux chamber with a diameter of 20 cm and a height of 11 cm was fastened on the PVC ring to create a headspace. The increase in CO_2_ and N_2_O concentrations in the headspace was determined after an incubation period of 10 minutes. The fluxes were determined assuming a linear increase over the incubation period. The validity of this assumption was checked by monitoring the concentration changes of one or two measuring points continuously over the incubation time.

### Statistical analysis

All statistical analyses were conducted in R version 4.3.0. A linear mixed model was used to determine the significance of differences in microbial gene abundance and chemical properties between treatments (control, P, K, PK, N, NPK, and PK+N) and seasons (spring and summer). The model was calculated with the “lme4” package (Bates et al., 2015) with treatment and season as fixed effects and time and site as random effects. Pairwise comparisons for fertilizer treatment were conducted with the “emmeans’’ package (Lenth et al., 2020). Redundancy analysis (RDA) was performed for microbial gene abundance and chemical properties in the “vegan” package (Dixon, 2003). The microbial genes and chemical properties listed in Tables 2 and 3 were included in the analysis. Correlations between the abundance of each gene and chemical properties were tested with Spearman’s coefficient analysis. Furthermore, the Random Forest was trained with chemical properties on the microbial gene abundance with the “randomForest ’’ package (Breiman, 2001). Predictions were made with a random forest of 1000 trees. The regression analysis between soil pH and gene abundance was conducted with the “lme4’’ package with time and site as random effects. We used MuMIn package (Barton, 2012) for calculating conditional R^2^ (Nakagawa and Schielzeth, 2013) . The regression model was selected from the linear model and binomial model based on the value of the akaike information criterion (AIC). The *P* and *R* values were calculated for the best-fit model and regression lines were shown only when the *P* value was significant (*P* < 0.05).

**Table 2.**
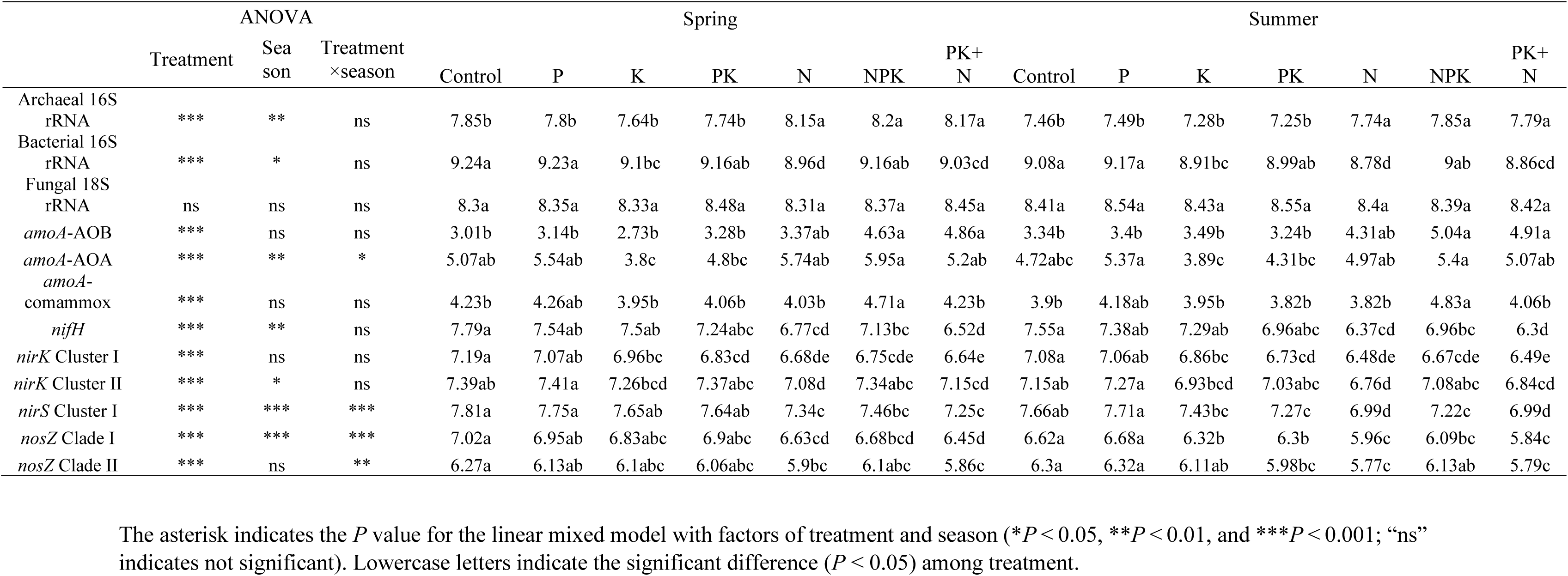
Microbial gene abundances in each season and treatment.

**Table 3.** Effect of fertilizer application on greenhouse gas emissions and soil chemical properties.

|  | ANOVA |  |  | Spring |  |  |  |  |  |  | Summer |  |  |  |  |  |  |
| --- | --- | --- | --- | --- | --- | --- | --- | --- | --- | --- | --- | --- | --- | --- | --- | --- | --- |
|  | Treat<br>ment | Sea<br>son | Treatmen<br>t×season | Control | P | K | PK | N | NPK | PK+N | Control | P | K | PK | N | NPK | PK+N |
| CO<sub>2</sub>-C<br>(kg/ha/day) | ns | ns | ns | 15.5a | 21.1a | 15.4a | 16.7a | 18.9a | 24.8a | 18.5a | 18.6a | 33.6a | 19.8a | 21a | 21.6a | 33.4a | 36.6a |
| N<sub>2</sub>O-N<br>(g/ha/day) | *** | ns | * | 0.517b | 0.642b | 0.693b | 0.527b | 3.95a | 3.82a | 3.17a | 0.357b | 0.353b | 0.448b | 0.502b | 1.81ab | 2.1a | 2.66a |
| pH (1:5<br>CaCl<sub>2</sub>) | *** | ns | * | 4.48ab | 4.56a | 4.35bc | 4.36abc | 4.09de | 4.23cd | 3.98e | 4.42ab | 4.53a | 4.35abc | 4.26bcd | 4.09de | 4.22cde | 4.05e |
| Moisture<br>(%) | ** | *** | * | 42.4a | 41.3a | 39.6ab | 37.1ab | 33b | 42.5a | 36.8ab | 24.9a | 27a | 23.5a | 24.4a | 21.5a | 26.7a | 23.4a |
| CN ratio | *** | ns | ns | 11ab | 10.6bc | 11.2a | 10.8ab | 10.3c | 11.1ab | 10.8bc | 11ab | 10.9bc | 11.3a | 11ab | 10.5c | 10.8ab | 10.7bc |
| NH<sub>4</sub><sup>+</sup>-N<br>(mg/kg) | *** | ns | *** | 11.3a | 11.4a | 10.9a | 9.65a | 11.9a | 21.6a | 8.98a | 10.6c | 10.7c | 11.7c | 12.5c | 27.1bc | 34.7ab | 49.4a |
| NO<sub>2</sub><sup>-</sup>-N<br>(mg/kg) | *** | ns | ns | 0.0848b | 0.14ab | 0.0827b | 0.0957b | 0.209ab | 0.257a | 0.168ab | 0.065b | 0.216ab | 0.0687b | 0.0853b | 0.18ab | 0.207a | 0.201ab |
| NO<sub>3</sub><sup>-</sup>-N<br>(mg/kg) | *** | ns | *** | 0.0833a | 0.45a | 0.142a | 0.0936a | 3.35a | 7.68a | 0.601a | 0.12b | 0.422b | 0.201b | 0.139b | 17a | 13.9a | 24.3a |
| K<sup>+</sup><br>(mmolc/kg) | *** | ns | * | 2.63b | 1.32b | 5.62a | 6.56a | 1.56b | 5.91a | 6.8a | 2.21c | 1.14cd | 5.94b | 7.38a | 0.825d | 6.32ab | 5.93b |
| P (resin)<br>(mg/kg) | *** | ns | ns | 30.4c | 145b | 20.2c | 178a | 33.3c | 139b | 143ab | 32.1c | 132b | 26.2c | 153a | 31c | 147b | 149ab |
| Ca<sup>2+</sup><br>(mmolc/kg) | *** | ns | * | 112b | 153a | 93b | 100b | 118b | 97.9b | 101b | 109b | 143a | 96.4b | 91.1b | 97.9b | 104b | 87.1b |
| Mg<sup>2+</sup><br>(mmolc/kg) | *** | ns | *** | 15.7b | 14.4b | 14b | 14.3b | 22.5a | 12.8b | 12.8b | 15.5ab | 15.1abc | 17.1a | 11.8c | 16.1ab | 13.2bc | 14.2abc |
| Cu (mg/kg) | *** | ns | ns | 1.34d | 7.86a | 9.15a | 5.88b | 7.11ab | 3.11cd | 5.28bc | 1.16d | 8.36a | 9.09a | 5.48b | 7.03ab | 3.53cd | 5.39bc |
| Fe (mg/kg) | *** | ns | ns | 365d | 983ab | 951ab | 930ab | 620cd | 1130a | 867bc | 365d | 955ab | 936ab | 956ab | 625cd | 1150a | 783bc |
| Mn (mg/kg) | *** | ns | ns | 8.71d | 27.1b | 38.5a | 21.8bc | 22bc | 10.3cd | 14.6bcd | 7.39d | 26.9b | 44.7a | 21.4bc | 21bc | 12.8cd | 19.2bcd |
| Zn (mg/kg) | *** | ns | ns | 1.92c | 9.58a | 9.82a | 8.25ab | 9.18a | 6.48b | 8.69a | 1.88c | 9.78a | 10a | 8.24ab | 9.72a | 6.64b | 9.54a |

The asterisk indicates the *P* value for the linear mixed model with factors of treatment and season (\**P* < 0.05, \*\**P* < 0.01, and \*\*\**P* < 0.001; “ns” indicates not significant). Values for the same property and treatment with different lowercase letters are significantly different (*P* < 0.05) between spring and summer.
† Carbon dioxide (CO<sub>2</sub>), nitrous oxide (N<sub>2</sub>O), ammonium (NH<sub>4</sub><sup>+</sup>), nitrite (NO<sub>2</sub><sup>-</sup>), nitrate (NO<sub>3</sub><sup>-</sup>), exchangeable potassium (K<sup>+</sup>), available phosphorus (P-resin), calcium (Ca<sup>2+</sup>) and magnesium (Mg<sup>2+</sup>), available iron (Fe), manganese (Mn), copper (Cu) and zinc (Zn).

## Result

### Abundance of N-cycle microbial genes

Two-way ANOVA analysis for the linear mixed model indicated a significant effect of fertilizer treatment on the genes of total archaea, total bacteria, *amoA*-AOB, *amoA*-AOA, *amoA*-comammox, *nifH*, *nirK*-I, *nirK*-II, *nosZ*-I, and *nosZ*-II (Table 2). Season variation influenced the abundance of total archaea, total bacteria, *amoA*-AOA, *nifH*, *nirK*-II, *nirS*-I, *nirS*-I, and *nosZ*-I. The application of N fertilizer had a negative effect on the abundance of *nirK*-I and *nirK*-II, *nirS*-I, *nosZ*-I, and *nosZ*-II genes while exerting a positive influence on *amoA*-AOA and *amoA*-AOB. The abundance of *amoA-*comammox was significantly higher in the NPK treatment than in other treatments.

### Effect of chemical properties on gene abundance

Two-way ANOVA analysis for the linear mixed model indicated that soil pH, moisture, soil mineral N (NH_4_⁺₋N, NO_2_ ^−^₋N, and NO_3_ ^−^₋N), available P, exchangeable cations (K^+^, Ca^2+^, and Mg^+^), cationic micronutrients [iron (Fe), manganese (Mn), copper (Cu) and zinc (Zn)] were significantly influenced by the fertilizer application (Table 3). Specifically, the soil pH was higher in order of P > Control > PK > K > NPK > N > PK+N treatment. Soil moisture was influenced by the interaction of fertilizer and season. In the summer season, the soil moisture decreased by 14.4% (averaged across all treatments) and there was no significant difference among the treatments.

The RDA analysis suggested the structure of the gene abundance differed in treatments with N, NPK, and PK+N compared to other treatments (Fig. 1a). Specifically, treatments containing N fertilizer show a higher abundance of *amoA*-AOB and *amoA*-AOA genes.

**Fig.1.**
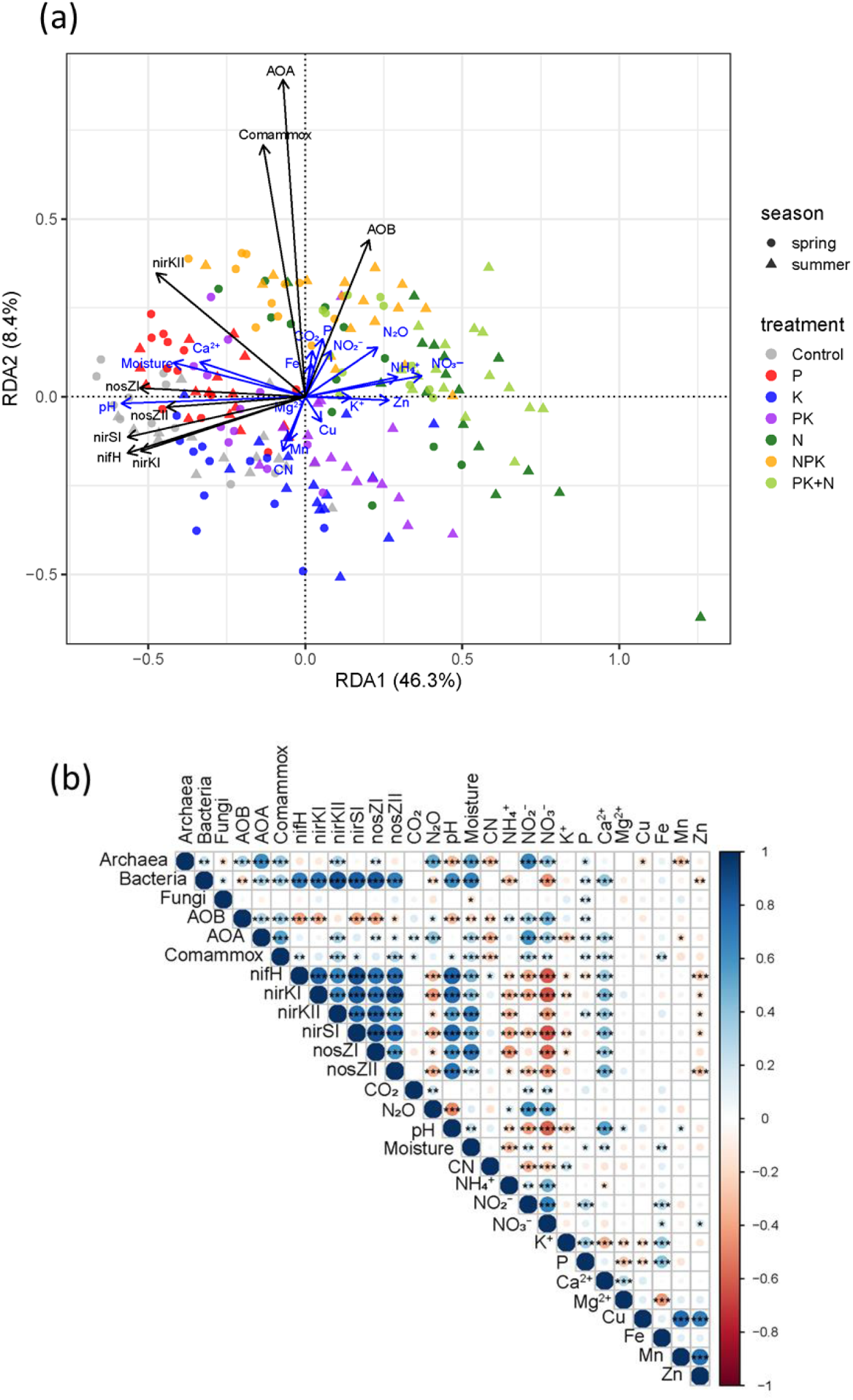
(a) Redundancy analysis (RDA) of the abundances of microbial genes and soil chemical properties. Only significant vectors (*P* < 0.05) for chemical properties are shown with blue arrows. (b) Heatmap of Spearman’s correlation coefficients between microbial gene abundances, soil chemical properties, and greenhouse gas emissions with *P* values (\**P* < 0.05, \*\**P* < 0.01, \*\*\**P* < 0.001).

Random Forest analysis showed that soil pH emerged as the most crucial factor positively correlating with the abundance of *nifH*, *amoA*-comammox, *nirK*-I, *nirS*-I, and *nosZ*-II (Fig. 2). The soil moisture was identified as the most influential factor positively correlated with *nirK*-II and *nosZ-*I. Further, regression analysis was performed with soil pH and quantified gene abundances. Soil pH was significantly positively correlated with the total bacteria, *amoA*-comammox, *nifH*, *nirK*-I and *nirK*-II, *nirS*-I, *nosZ*-I, and *nosZ-*II.

**Fig. 2.**
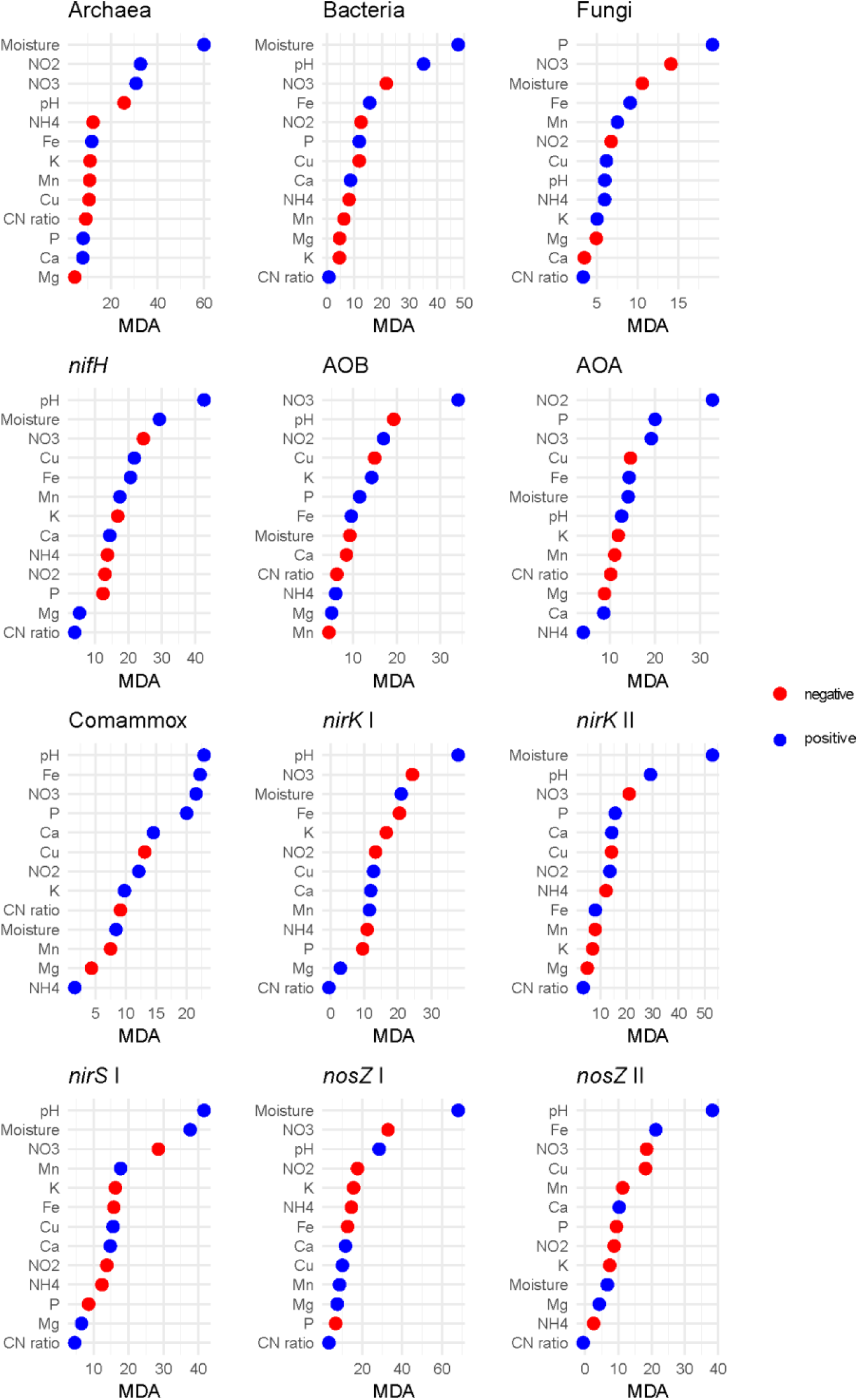
Mean decrease accuracy (MDA) for Random Forest analysis for chemical properties and gene abundance. The color of the plot indicates blue; positive or red; negative correlation with each chemical factors.

### CO2 and N2O emissions

The CO_2_ emissions were not significantly different among treatments while N_2_O emissions were significantly higher when mineral N was added (N, NPK, PK+N) in both spring and summer (Table 3). No significant differences were observed in N_2_O emissions among the N, NPK, and PK+N treatments. The N_2_O emission was similar among the treatments without N fertilizer (Control, P, K, PK). The emission of N_2_O was significantly positively correlated with the gene abundance of total archaea and *amoA*-AOA. Conversely, N_2_O emission showed a significant negative correlation with the abundance of genes related to total bacteria, *nifH*, *nirK*-I and *nirK-*II, *nirS*-I, and *nosZ*-I and *nosZ-*II (Fig. 1b).

## Discussion

In the present study, the shift of microbial N-cycling genes, such as *nifH*, *nirS*-I, *nirK*-I, and *nosZ*-II, is more effectively explained by the change of soil pH induced by long-term chemical fertilizer application than by the short-term nutrient inputs of fertilizer applications and availability of nutrients, such as mineral N and available P, provide by fertilization. This is also clear from the fact that in the soils that receive similar nutrients (e.g., NPK vs. PK+N), the abundance of microbial N-cycling genes separated based on the soil pH (Fig. 3). Soil pH is often described as one of the most important factors shaping microbial diversity and functionally. In general, the acidification of the soil pH is associated with a decrease in the abundance and diversity of the microbial community (Dong et al., 2022; Griffiths et al., 2011; Rousk et al., 2010). However, it remained unclear whether soil pH or nutrient availability played a more significant role in microbial growth in the acidic grassland. Previous global meta-analyses of grasslands have reported a positive effect of N application on the gene abundance of *nirK*, *nirS*, and *nosZ* (You et al., 2022). Conversely, in the present study, long-term N application was associated with a reduction in denitrifiers concurrent with a decrease in soil pH. Similarly, for diazotrophs previous studies suggest that the application of PK fertilizer increases the abundance of the *nifH* gene in the cropland with soil pH around 5 to 6 (Hu et al., 2019; Wang et al., 2017), whereas our study demonstrated the negative effect of PK fertilizer on the *nifH* gene. Overall, the findings suggest that the decrease in soil pH resulting from long-term fertilizer application overrides the effects of nutrient availability from fertilizers on diazotrophs and denitrifiers in acidic grasslands.

**Fig. 3.**
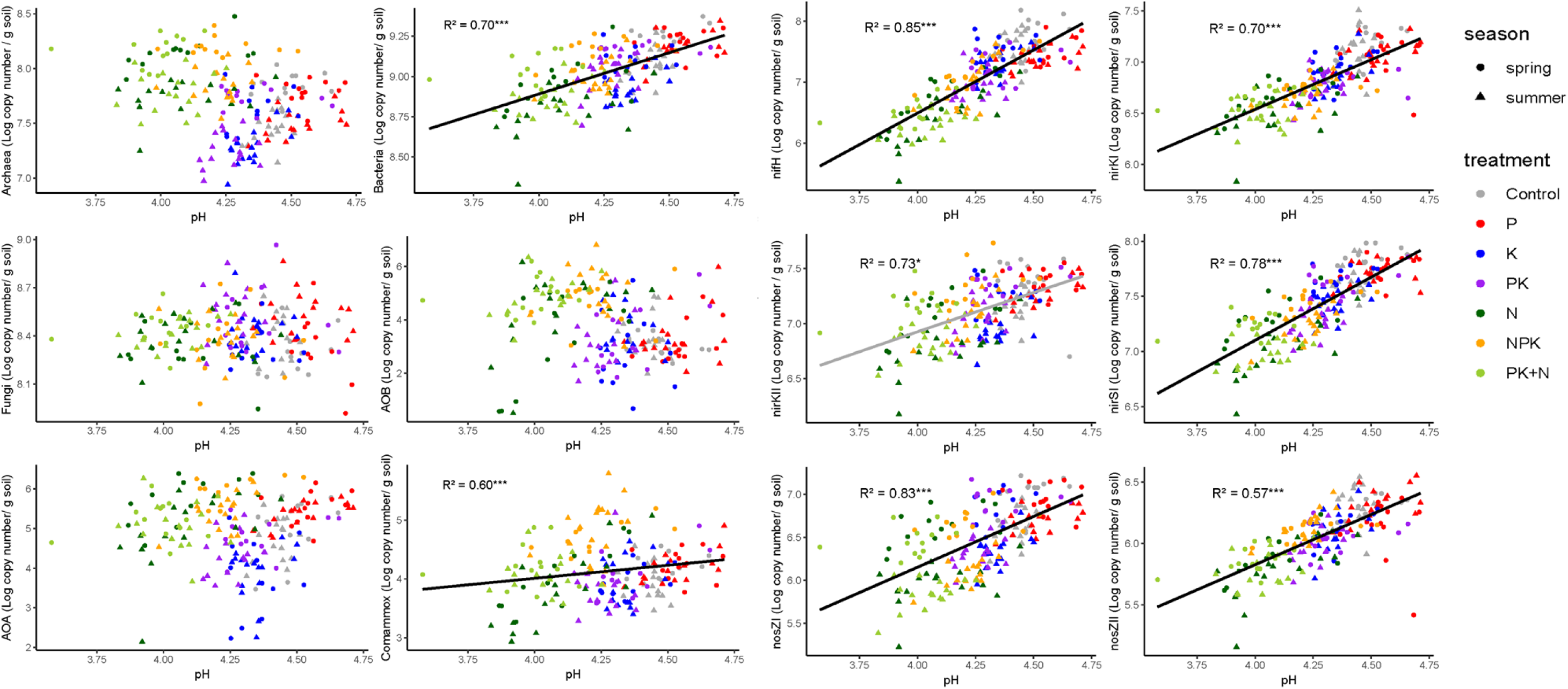
Microbial gene abundance was plotted against soil pH. The best regression model was selected and from linear (black) or polynomial (grey) model on the basis of the Akaike information criterion (AIC). The regression lines were shown only when the *P* value was significant (*P* < 0.05). Conditional R^2^ values are shown with *P* values (\**P* < 0.05, \*\**P* < 0.01, \*\*\**P* < 0.001).

The shift in soil pH caused by long-term fertilizer application was dependent on the type of fertilizer and their combination. Only the application of single P (superphosphate) increased soil pH, whereas when applied together with potassium K (PK), the soil pH was lower than in the control. The effect of superphosphate on soil pH is reported to be controversial (Sarangi and Jena, 2020; Williams and Donald, 1957). However, a recent study indicates that phosphate could increase the negative charge on the soil surface, especially at low soil pH, leading to greater soil adsorption of hydrogen ions (H^+^) and mitigation of soil acidity (Barrow et al., 2022). The increase in potassium ions (K^+^) with the application of K fertilizer might compete with H^+^ in the negative charges of the soil, thus causing a decrease in soil pH when PK was applied. Meanwhile, the application of N fertilizer reduced soil pH because the oxidation process of ammonium (NH_4_ ^+^) to nitrate (NO_3_ ^−^) produces H^+^. When PK was applied simultaneously with N (NPK), the reduction of soil pH was mitigated compared to a single N application. However, when they were separately applied (PK+N), the soil pH dropped to the same level as with a single N application. Previous studies indicate that N fertilizer accelerates soil acidification more than NPK fertilizer in acidic soil (Daba et al., 2021; Tuyen et al., 2006). However, our results from PK+N imply that rather than the presence of PK, the single application of N could be the main reason for the decrease in soil pH. Further study is necessary to explain the mechanisms behind this complex chemical interplay including the association with plants.

In addition to soil pH, soil moisture content was also a crucial factor influencing the temporal shift in denitrifier gene abundance (Fig. 2). Specifically, the abundance of *nirK*-II and *nosZ*-I showed a higher response to soil moisture compared to other genes related to denitrification. This finding aligns with a previous study indicating that *nosZ*-I was more sensitive to soil moisture than *nosZ*-II (Xu et al., 2020). Furthermore, transcript analysis revealed transcripts of *nirK*-II and *nosZ*-I were found to increase simultaneously (Wei et al., 2021). This suggests that microbes possessing these genes share a similar ecological niche. *nirK*-II derives from diverse microbes, including Alpha-, Beta-, Gamma-, Delta- and Epsilon-*proteobacteria*, as well as *Bacteroidetes*, *Chloroflexi*, and *Spirochetes* (Wei et al., 2015). In contrast, *nosZ*-I derives from Alpha-, Beta-, and Gamma-*proteobacteria* (Jones et al., 2013). The overlap of the phylogeny of microbes possessing those genes needs to be checked in future studies.

Compared to denitrifying microbes, nitrifying microbes, including *amoA*-AOA and *amoA*-AOB showed no significant correlation with soil pH, while *amoA*-comammox exhibited positive correlation. Instead, the abundance of *amoA*-AOB and *amoA*-AOA increased with the application of N, NPK, and PK+N, and the abundance of *amoA*-comammox was significantly more abundant with NPK treatment (Table 2). This is likely because ammonia-oxidizers are autotrophic microbes that directly rely on N and are sensitive to the temporal increase of N availability (Mitsuta et al., 2023), whereas denitrifiers are heterotrophic microbes that may also depend on other nutrients (Sun and Jiang, 2022). It is essential to note that the abundance of AOA was also sensitive to P and K fertilizer. The sole application of P had a positive impact on the abundance of AOA. This finding aligns with a previous study indicating that P application is favored by AOA rather than AOB in acidic forest soil (Tang et al., 2016). P fertilizer application could facilitate organic matter decomposition and N mineralization, leading to increased available N for AOA. Meanwhile, the abundance of AOA decreased with PK or K application. One possible explanation is that K application negatively affects NH_4_⁺ availability as K^+^ replaces the NH_4_⁺ on soil negative charges (Nieder et al., 2011). However, we did not find any negative effect on NH_4_⁺-N concentration in the soil. Similarly, the inhibition effect of K^+^ on nitrification has been reported by previous studies (Golden et al., 1981; Li et al., 2020) but the mechanisms behind this inhibitory effect have not been investigated so far. For comammox, only the application of NPK had a positive effect on its abundance, whereas the application of N and PK+N had a negative effect. This is likely because NPK treatment had higher soil pH than N and PK+N treatment, indicating that comammox is sensitive to both soil pH and N availability when soil pH is lower than around 4.7. This is in line with previous studies that found comammox to be positively correlated with both soil pH and NH_4_⁺ availability (Osburn and Barrett, 2020; Sun et al., 2021). Overall, the type of chemical fertilizer appears to be more critical for nitrifier abundance than for denitrifiers in acidic grasslands.

The N_2_O emission was increased with the application of N, NPK, and PK+N treatments compared to other treatments (Table 3). There was no significant difference in the emission among these N treatments. The emission was significantly positively correlated with the abundance of total archaea and *amoA*-AOA gene abundance, whereas it negatively correlated with *nirK*, *nirS*, and *nosZ* abundance as those gene abundances were decreased with the application of N fertilizer (Fig. 1b and Table 2). Although we observed a tendency for the ratio of *nirK* and *nirS* to *nosZ* to increase with the application of N there was no direct correlation with N_2_O emissions (Fig S1 and S2). This lack of correlation may be attributed to the stable abundance of denitrifiers during the sampling time points, while N_2_O emissions exhibited a temporal increase corresponding to precipitation events (Fig. S3 and S4). Notably, denitrifiers harbor the *nirK*, *nirS*, and *nosZ* mRNA corresponding to the N fertilizer application and decline of the oxygen level after rainfall event (Uchida et al., 2014), thus transcriptome analysis remains a subject for further study in this field. However, our result still yields valuable insight into how the long-term effect of chemical fertilizer influences the absolute gene abundances associated with N_2_O emission, particularly revealing the unique patterns in the acidic grassland.

## Conclusion

The long-term application of synthetic fertilizer to acidic grassland leads to a specific distribution of the N-cycling microbial community. Our study demonstrated that the difference in soil pH shaped by the long-term application of fertilizer overrides the effect of nutrient availability from fertilizer application on diazotrophs and denitrifiers abundance. On the contrary, the abundance of nitrifiers increased with the application of N fertilizer, despite the decrease of the soil pH due to the application of N. These findings provide crucial insights into how the long-term management of fertilizer affects microbial N-cycling, emphasizing pH as the most influential factor shaping microbial genes essential for environmental processes.

## Acknowledgments

The authors thank Ji Li, Jingjing Chang, and Shuaimin Chen for their assistance with field sampling, Gert van Dorland for the execution of the trace gas measurements, Agata S. Pijl for molecular laboratory technical assistance, and Dr. Heitor Cantarella for the nutrient analysis at the Agronomical Institute of Campinas—IAC (Brazil). We thank René Schils for the coordination of KB36 Biodiversity in a Nature Inclusive Society project.

