## Supplementary material for "Unveiling the dominant role of soil pH in shaping nitrogen cycling microbial gene abundances: Insights from 65-years of chemical fertilizer selection in acidic grassland meadow": Table S1

**Cloning for qPCR standards**

For each gene PCR amplicon was prepared using microorganisms containing the gene. The PCR products were purified using QIAquick® PCR purification kit (Qiagen, Germany) according to the manufacturer's protocol. Ligation of the purified PCR products into the plasmid vector was performed using pGEM®-T Easy Vector (Promega, USA) then transformed into JM109 Competent Cells High Efficiency (Promega,USA) according to the manufacturer’s protocol. After 1.5-hour incubation at 37 degrees in incubator shaker the cell suspension was plated onto the agar plate. After overnight incubation of the plate a single white colony was picked up and transferred to lysogeny broth media containing the antibiotic ampicillin and incubated overnight. Plasmids were extracted from the cultures using Qiaprep Spin Miniprep Kit V 1.0 (Qiagen, Germany). The concentration and quality of the plasmids were checked using the NanoDrop ND-1000 spectrophotometer (NanoDrop Technologies, DE, USA). Standard curves generated with a series of 1:10 dilutions of the plasmids. The efficiencies of the standards were described in the Table S1 and *R* values were above 0.98 among all standards. For the genes *nosZ* clade I and *nirK* cluster I the standard efficiency was low due to the adjustment of the primer concentration and annealing temperature to eliminate the smear generation based on the observation with electrophoresis.

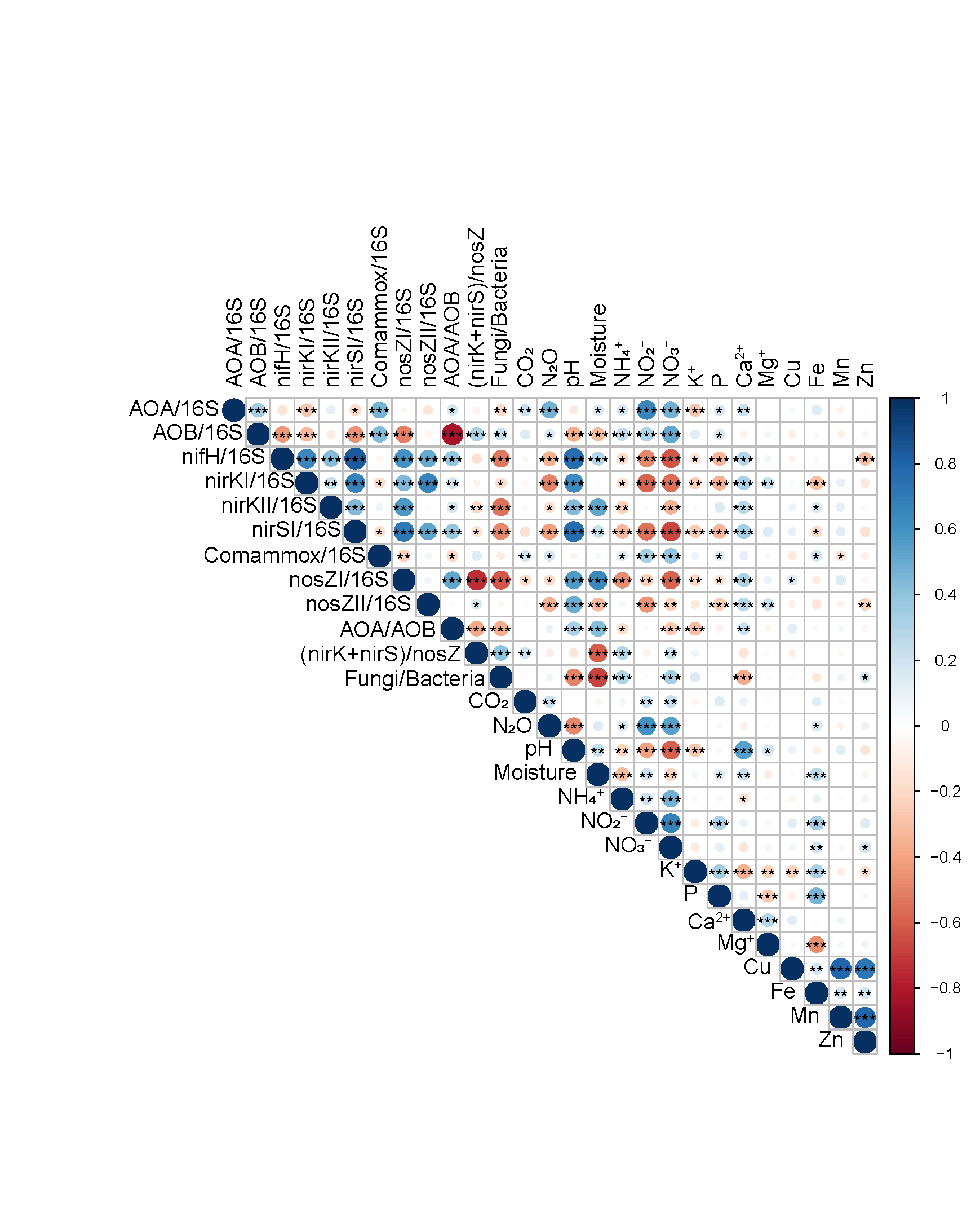

Fig.S1. Heatmap of Spearman's correlation coefficients for relative microbial N-cycling gene abundances to prokaryotes abundance, and ratio of *nirK*+*nirS* to *nosZ* genes, *amoA*-AOB to *amoA*-AOA, Fungi to bacteria, soil chemical properties, and greenhouse gas emissions with *P* values (**P* < 0.05, ***P* < 0.01, ****P* < 0.001).

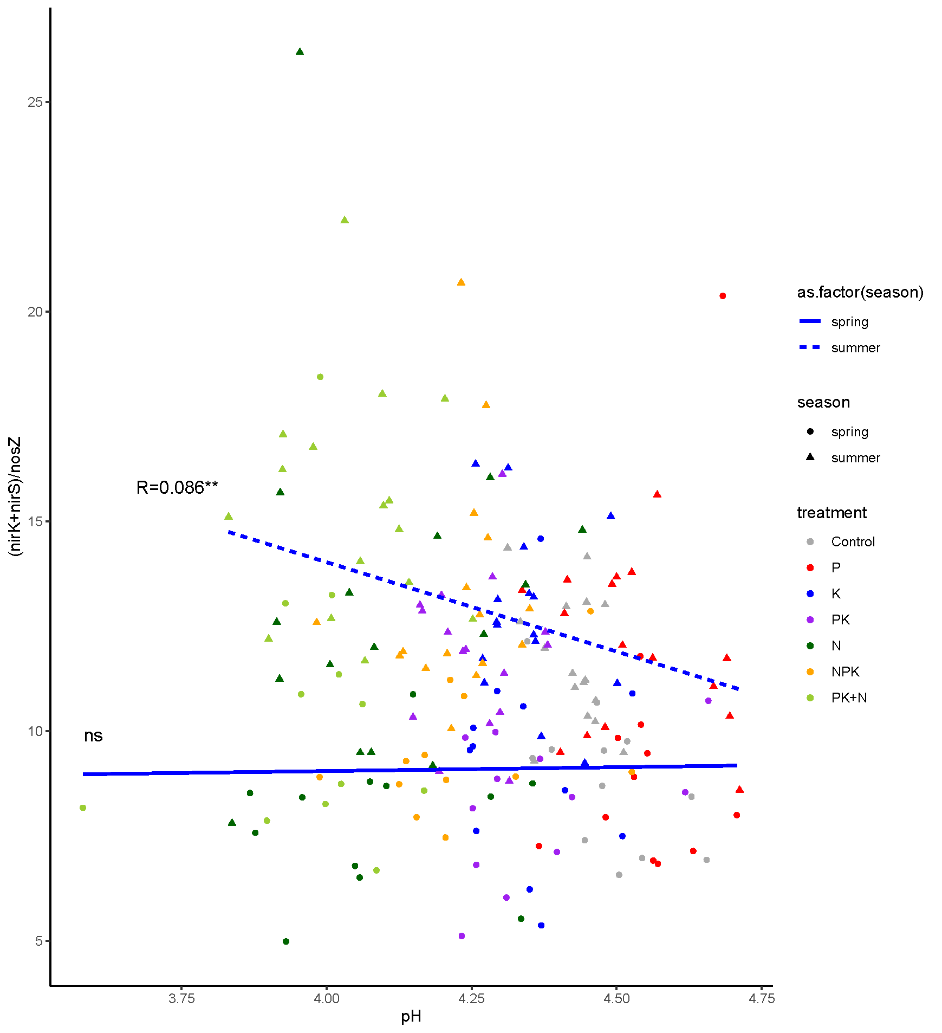

Fig. S2. Relationship of soil pH and (nirK+nirS)/nosZ at each season.

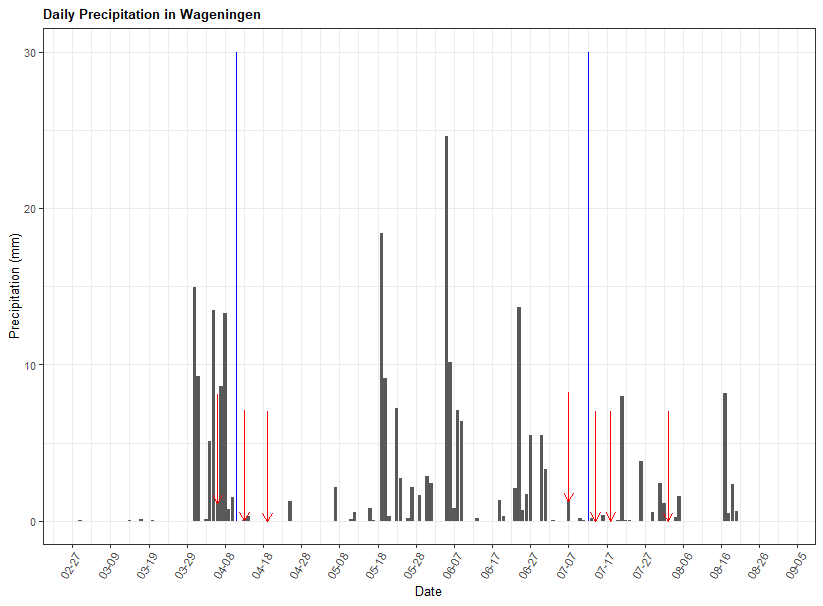

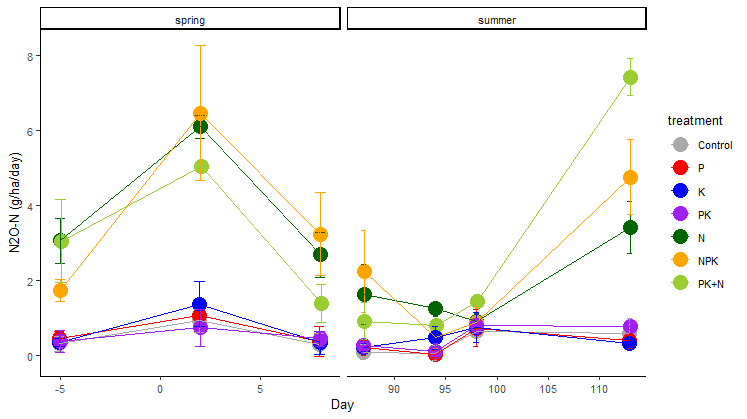

Fig. S3. Precipitation in Wageningen and N_2_O emission. In the precipitation the red arrow indicates the date of sampling. Blue line indicates the fertilizer application. Precipitation data was obtained from open-source Visual Crossing website (https://www.visualcrossing.com).

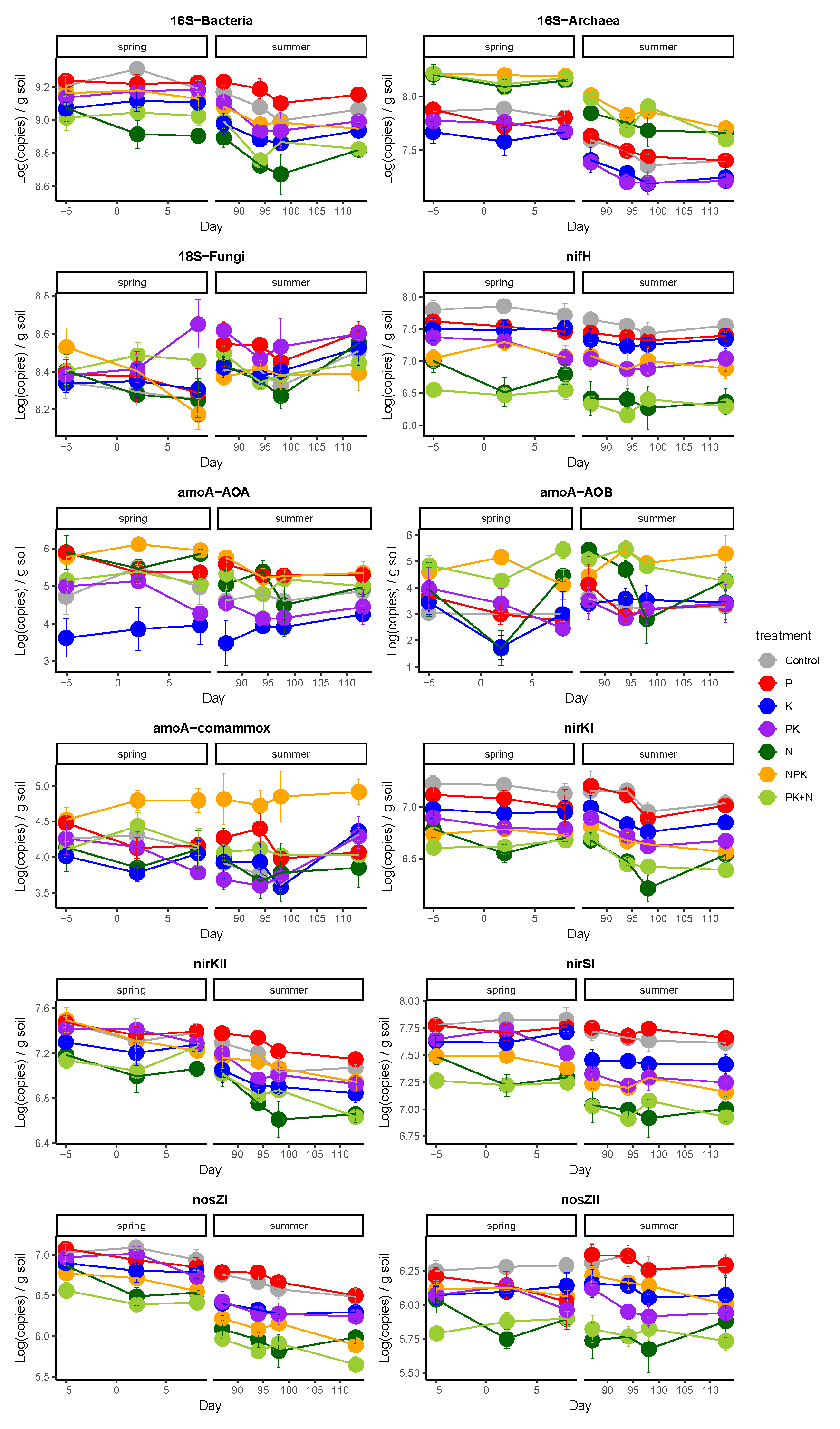

Fig.S4. Microbial gene abundance with seven timepoints. The error bar shows standard error.

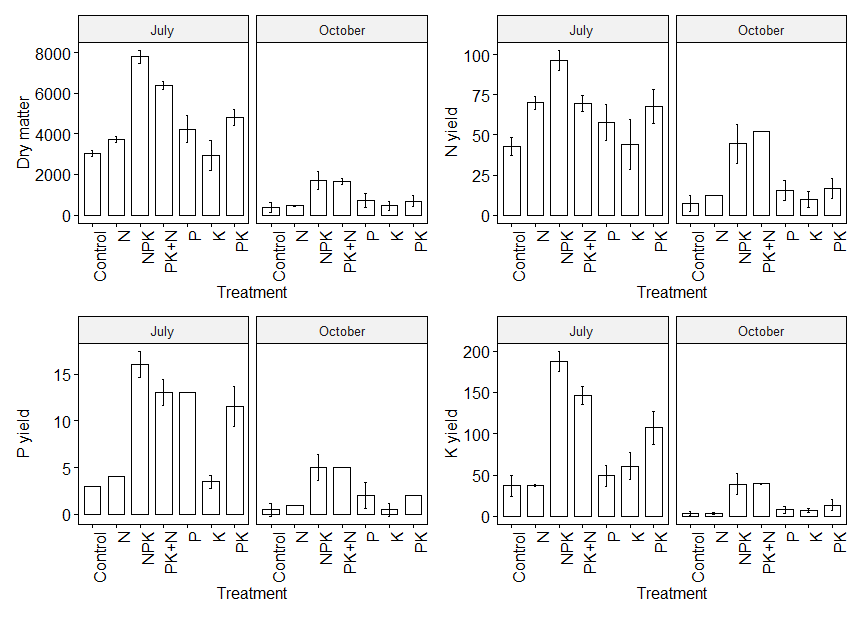

Fig. S5. Plant yield on first and second harvest. The error bar shows standard error.

Table S1. Sequences of the primers and the optimal PCR conditions.

| Target gene | Primers | Primer Sequence | Reaction | Cycling conditions | Standard efficiency (%) | Reference |
| --- | --- | --- | --- | --- | --- | --- |
| Archaeal 16S rRNA | ARC915F | 5’-AGGAATTGGCGGGGGAGCAC-3’ | 6 μL of Sybrgreen Bioline SensiFAST SYBR non-rox mix, 0.48 μL of each primer (10 pmol), and 2 μL of DNA (2.5 ng) | 95°C-3 min.; 40x 95°C-5s, 60°C-10s, 72°C-20s | 89.7 | Klindworth et al. (2013) |
|  | 1017R | 5’-GGCCATGCACCWCCTCTC-3’ |  |  |  |  |
| Bacterial 16S rRNA | EUB338 | 5’-ACTCCTACGGGAGGCAGCAG-3’ | 6 μL of 2 SensiMix SYBR no-rox kit, 0.125 μL of each primer (10 pmol), 0.3 μL of BSA, and 2 μL of DNA (1 ng) | 95°C-10 min.; 40x 95°C-15s, 60°C-15s, 72°C-20s | 94.7 | Fierer et al. (2005) |
|  | EUB518 | 5’-ATTACCGCGGCTGCTGG-3’ |  |  |  |  |
| Fungal 18S rRNA | FF390 | 5'-CGATAACGAACGAGACCT-3' | 6 μL of Sybrgreen Bioline SensiFAST SYBR non-rox mix, 0.25 μL of each primer (10 pmol), 1.25 μL of BSA, and 2 μL of DNA (1 ng) | 95°C-2 min.; 40x 95°C-5s, 52°C-10s, 72°C-20s | 93 | Vainio and Hantula (2000) |
|  | FR1 | 5'-AICCATTCAATCGGTAIT-3' |  |  |  |  |
| *amoA*-AOB | amoA1F | 5’-GGGGTTTCTACTGGTGGT-3’ | 6 μL of Sybrgreen Bioline SensiFAST SYBR non-rox mix, 0.125 μL of each primer (10 pmol), and 4 μL of DNA (10 ng) | 95°C-10 min.; 45x 95°C-10s, 64°C-25s | 84.9 | Rotthauwe et al. (1997) |
|  | amoA2R | 5’-CCCCTCKGSAAAGCCTTCTTC-3’ |  |  |  |  |
| *amoA*-AOA | Arch-amoAF | 5’-STAATGGTCTGGCTTAGACG-3’ | 6 μL of Sybrgreen Bioline SensiFAST SYBR non-rox mix, 0.125 μL of each primer (10 pmol), and 4 μL of DNA (10 ng) | 95°C-5 min.; 45x 95°C-10s, 64°C-10s, 72°C-20s | 89.9 | Francis et al. (2005) |
|  | Arch-amoAR | 5’-GCGGCCATCCATCTGTATGT-3’ |  |  |  |  |
| *amoA*-comammox | Ntsp-amoA 162F | 5’-GGATTTCTGGNTSGATTGGA-3’ | 6 μL of Sybrgreen Bioline SensiFAST SYBR non-rox mix, 0.48 μL of each primer (10 pmol), and 4 μL of DNA (1 ng) | 95°C-3 min.; 45x 95°C-10s, 63°C-10s, 72°C-20s | 77.3 | Fowler et al. (2018) |
|  | Ntsp-amoA 359R | 5’-WAGTTNGACCACCASTACCA-3’ |  |  |  |  |
| *nifH* | PolF | 5’-TGCGAYCCSAARGCBGACTC-3’ | 6 μL of Sybrgreen Bioline SensiFAST SYBR non-rox mix, 0.48 μL of each primer (10 pmol), and 2 μL of DNA (5 ng) | 95°C-3 min.; 45x 95°C-5s, 65°C-10s, 72°C-20s | 92.1 | Poly et al. (2001) |
|  | PolR | 5’-ATSGCCATCATYTCRCCGGA-3’ |  |  |  |  |
| *nirK* Cluster I | NirKC1F | 5’-ATGGCGCCATCatggtnytncc-3’ | 6 μL of Sybrgreen Bioline SensiFAST SYBR non-rox mix, 0.2 μL of each primer (10 pmol), and 4 μL of DNA (1 ng) | 95°C-3 min.; 45x 95°C-10s, 58°C-10s, 72°C-30s | 65.8 | Wei et al. (2015) |
|  | NirKC1R | 5’-TCGAAGGCCTCGatnarrttrtg-3’ |  |  |  |  |
| *nirK* Cluster II | nirKC2F | 5’-TGCACATCGCCAACggnatgtwygg-3’ | 6 μL of Sybrgreen Bioline SensiFAST SYBR non-rox mix, 0.48 μL of each primer (10 pmol), and 2 μL of DNA (2.5 ng) | 95°C-3 min.; 45x 95°C-5s, 58°C-10s, 72°C-20s | 71.3 | Wei et al. (2015) |
|  | nirKC2R | 5’-GGCGCGGAAGATGshrtgrtcnac-3’ |  |  |  |  |
| *nirS* Cluster I | nirSC1F | 5’-ATCGTCAACGTCaargaracvgg-3’ | 6 μL of Sybrgreen Bioline SensiFAST SYBR non-rox mix, 0.48 μL of each primer (10 pmol), and 4 μL of DNA (1 ng) | 95°C-3 min.; 40x 95°C-10s, 58°C-10s, 72°C-30s | 78.4 | Wei et al. (2015) |
|  | nirSC1R | 5’-TTCGGGTGCGTCttsabgaasag-3’ |  |  |  |  |
| *nosZ* Clade I | nosZ2F | 5’-CGCRACGGCAASAAGGTSMSSGT-3’ | 6 μL of Sybrgreen Bioline SensiFAST SYBR non-rox mix, 0.125 μL of each primer (10 pmol), 0.3 μL of BSA, and 4 μL of DNA (1 ng) | 95°C-5 min.; 40x 95°C-10s, 64°C-10s, 72°C-20s | 62.8 | Henry et al. (2006) |
|  | nosZ2R | 5’-CAKRTGCAKSGCRTGGCAGAA-3’ |  |  |  |  |
| *nosZ* Clade II | nosZII-F | 5’-CTIGGICCIYTKCAYAC-3’ | 6 μL of Sybrgreen Bioline SensiFAST SYBR non-rox mix, 0.25 μL of each primer (10 pmol), and 4 μL of DNA (1 ng) | 95°C-3 min.; 45x 95°C-20s, 60°C-60s, 72°C-30s | 77.1 | Jones et al. (2013) |
|  | nosZII-R | 5’-GCIGARCARAAITCBGTRC-3’ |  |  |  |  |
